# A comparison of bivariate frequency domain measures of electrophysiological connectivity

**DOI:** 10.1101/459503

**Authors:** Roberto D. Pascual-Marqui, Pascal Faber, Toshihiko Kinoshita, Kieko Kochi, Patricia Milz, Nishida Keiichiro, Masafumi Yoshimura

## Abstract

The problem of interest here concerns electrophysiological signals from two cortical sites, acquired as invasive intracranial recordings, or from non-invasive estimates of cortical electric neuronal activity computed from EEG or MEG recordings (see e.g. https://doi.org/10.1101/269753). In the absence of other sources, these measured signals consist of an instantaneous linear mixture of the true, actual, unobserved local signals, due to low spatial resolution and volume conduction. A connectivity measure is unreliable as a true indicator of electrophysiological connectivity if it is not invariant to mixing, or if it reports a significant connection for a mixture of independent signals. In (Vinck et al 2011 Neuroimage 55:1548) it was shown that coherence, imaginary coherence, and phase locking value are not invariant to mixing, while the phase lag index (PLI) and the weighted version (wPLI) are invariant to mixing. Here we show that the lagged coherence (LagCoh) measure (2007, https://arxiv.org/abs/0711.1455), not studied in Vinck et al, is invariant to mixing. Additionally, we present here a new mixture-invariant connectivity statistic: the “standardized imaginary covariance” (sImCov). We also include in the comparisons the directed PLI (dPLI) by Stam et al (2012 Neuroimage 62:1415). Fourier coefficients for “N” trials are generated from a linear unidirectional causal time domain model with electrophysiological delay “k” and regression coefficient “b”. 1000 random data sets of “N” trials are simulated, and for each one, and for each connectivity measure, non-parametric randomization tests are performed. The “true positive detection rate” is calculated as the fraction of 1000 cases that have significant connectivity at p<0.05, 0.1, and 0.2. The connectivity methods were compared in terms of detection rates, under non-mixed and mixed conditions, for small and large sample sizes “N”, with and without jitter for “k”, and for different values of signal to noise. Under mixing, the results show that LagCoh outperforms wPLI, PLI, dPLI, and sImCov. Without mixing, LagCoh and sImCov outperform wPLI, PLI, and dPLI. Finally, it is shown that dPLI is an invalid estimator of flow direction, i.e. it reverses and “goes against the flow” by merely changing the sign of one of the time series, a fact that violates the basic definition of Granger causality. For the sake of reproducible research, the supplementary material includes Delphi Pascal source code and all detailed result files in human readable format.

## 2. Introduction

The problem of interest here concerns electrophysiological signals from two cortical sites, obtained as invasive intracranial recordings, or from non-invasive estimates of cortical electric neuronal activity computed from EEG or MEG recordings by means of an inverse solution (see e.g. Pascual-Marqui et al 2018 for a comparative review of linear inverse solutions).

Consider a hypothetical ideal situation where there are no other sources present. In this case, these measured signals consist of an instantaneous linear mixture of the true, actual, unobserved local signals, due to low spatial resolution and volume conduction. Thus, the observed signals are artifactually “similar”, and conventional measures of electrophysiological connectivity, such as coherence and phase locking value, can produce significant false positive results.

A general solution to the mixing problem, denoted “innovations orthogonalization”, was given in Pascual-Marqui et al (2017). It provides an appropriate “unmixing” for the general multivariate time series case. After unmixing, any connectivity measure can be calculated, even measures such as coherence and phase locking value.

An alternative solution to this problem is to find bivariate measures of electrophysiological connectivity that are invariant to mixing. In this sense, a connectivity measure is unreliable as a true indicator of electrophysiological connectivity if it is not invariant to mixing, or if it reports a significant connection for a mixture of independent signals.

In the study by Vinck et al (2011), it was shown that coherence (see e.g. Brillinger 2001), imaginary coherence (Nolte et al 2004), and phase locking value (Lachaux et al 1999) are not invariant to mixing, while the phase lag index (PLI) (Stam et al 2007) and the weighted version (wPLI) (Vinck et al 2011) are invariant to mixing.

In this study we show that the lagged coherence measure is invariant to mixing. The lagged coherence can be found in (Eq. 22 in Pascual-Marqui 2007a; Eqs. 29 and 31 in Pascual-Marqui 2007b; Eqs. 3.13, 3.14, and 3.17 in Pascual-Marqui et al 2011b). It was not studied in (Vinck et al 2011).

Additionally, we present here a new mixture-invariant connectivity statistic: the “standardized imaginary covariance” (sImCov).

We also include in the comparisons the directed PLI (dPLI) by Stam and van Straaten (2012).

A good measure of electrophysiological connectivity must minimally possess the following properties:

1. It must be mixing-invariant, i.e., invariant to instantaneous linear mixtures. Therefore, it must produce no false positive connections for any mixture of independent signals.
2. It must detect the highest possible rate of true positive connections in the presence of connections with electrophysiological time delays.

The comparison of connectivity measures is based on quantifying these two properties under simulations. An outline of the procedure follows:

1. Fourier coefficients at a selected frequency are generated for two signals satisfying a unidirectional causal time domain model with electrophysiological time delay. The number of Fourier coefficients corresponds to the sample size, which is equal to the number of trials or epochs. Jitter and additive measurement noise are included.
2. Thus, two sets of data are available: the original noisy non-mixed signals, and their mixed version. Mixing is implemented with a positive definite 2×2 matrix of mixing coefficients.
3. For the non-mixed and for the mixed signals, each connectivity measure is computed, and non-parametric randomization significance tests are carried out.
4. Steps (1), (2), and (3) are repeated for 1000 random realizations, and the number of true positive detections (at p<0.05, p<0.1, and p<0.2) in 1000 experiments are counted. These counts correspond to the rate of true positive connections detected, and constitutes the basis for ranking the connectivity measures.

All steps above (1) through (4), are carried out for small and large sample sizes, with and without jitter, and for different values of signal to noise. A non-exhaustive total of 19 combinations of sample size, noise, and jitter were computed, allowing a ranking of connectivity measures.

The results show that under mixing, the lagged coherence measure outperforms wPLI, PLI, dPLI, and sImCov. For the ideal, non-mixed signals case, lagged coherence and sImCov outperform wPLI, PLI, and dPLI.

Finally, it is shown that dPLI is an invalid estimator of flow direction, i.e. it reverses and “goes against the flow” by merely changing the sign of one of the time series, a fact that violates the basic definition of Granger causality. This is particularly relevant, because the sign of electrophysiological signals are irrelevant in many instances, as explained later in more detail in the discussion.

## 3. Notation and basic definitions for bivariate time series and positive definite mixtures

Let *u_tk_* and *v_tk_* denote real-valued time series, representing the true, unobserved, unmixed cortical electric neuronal activity at two sites. There are available “N” trials or epochs, for *k* = 1…*N*, with each epoch consisting of *N_T_* discrete time samples, for *t =*1…*N_T_*. The complex valued discrete Fourier transform coefficients are denoted as *u*_ω*k*_ and *v*_ω*k*_, for discrete frequency ω.

Define the complex vector:

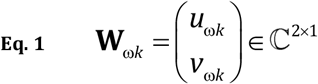

with Hermitian variance-covariance matrix:

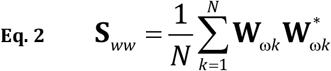

where the superscript “*” denotes, in general, transpose and complex conjugate.

The corresponding observed, recorded time series are denoted as *x_tk_* and *y_tk_*, with Fourier coefficients *x*_ω*k*_ and *y*_ω*k*_, for which the following vector is defined:

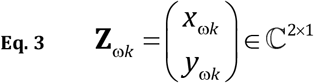

This vector is a linear, instantaneous mixture of the true, unobserved vector, i.e.:

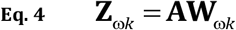

where the matrix 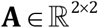 in this study is assumed to be positive definite.

Note that in the case of signals of estimated cortical current density based on a linear inverse solution for EEG/MEG (such as those reviewed and compared in Pascual-Marqui et al 2018), the mixing matrix **A** is a submatrix of the resolution matrix, which is non-negative definite. Some technical details about the resolution matrix and its properties can be found in e.g. Pascual-Marqui et al (2011a).

The Hermitian variance-covariance matrix for the observations **Z** is:

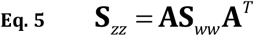

Let:

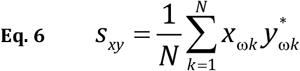

denote the Hermitian complex-valued covariance between any two variables “x” and “y”.

The relation between the imaginary part of the mixed covariance and the imaginary part of the true unmixed covariance can be found in Vinck et al (2011). It can be derived from Eq. 5, which gives:

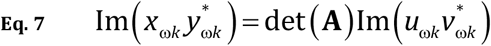

and:

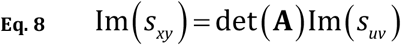

where:

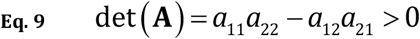

is the determinant of the mixing matrix, which is assumed to be positive in this study. In Eq. 7 and Eq. 8, Im (•) returns the imaginary part of a complex valued argument.

## 4. Some bivariate frequency domain measures of electrophysiological connectivity

### 4.1. >The imaginary part of the complex-valued coherence (ImCoh)

The imaginary part of the complex-valued coherence was introduced by Nolte et al (2004). This was probably the first study to address the problem of artifactual connections inferred from the classical coherence measure, due to volume conduction. The reasoning for using the imaginary part was that since it depends on time delays, it should not be affected by instantaneous volume conduction. However, as shown in Vinck et al (2011), ImCoh in Eq. 10 is not invariant to mixing, i.e. it is not invariant to volume conduction. Despite this fact, the imaginary coherence will be considered in this study, because it is still of great interest to see how its performance compares to other measures in the ideal case of no-mixing.

The full complex valued coherence is:

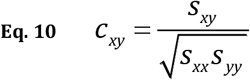

From here, the imaginary part, is denoted as:

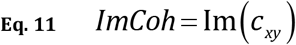

### 4.2. The phase lag index (PLI)

The phase lag index (PLI) was introduced by Stam et al (2007). It is defined as:

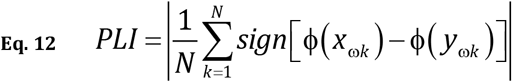

where Φ(•) returns the real valued phase angle of the complex valued argument defined for the range[(−π)···(+π)],and where, for any real-valued *x*:

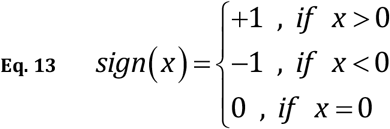

An equivalent definition for the PLI is:

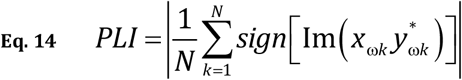

See e.g. Eq. 7 in Vinck et al (2011).

A non-zero PLI indicates that there is lagged synchronization, and it was shown to be mixing-invariant in Vinck et al (2011). This can be seen by plugging Eq. 7 into Eq. 14.

### 4.3. The weighted phase lag index (wPLI)

The weighted phase lag index (wPLI) was introduced by Vinck et al (2011). It is defined as:

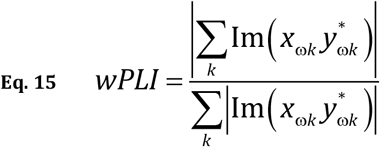

(Vinck et al (2011) showed that this measure is mixing-invariant (by plugging Eq. 7 into Eq. 15), and also showed that it offers several advantages over the original PLI.

### 4.4. The directed PLI (dPLI)

The directed PLI (dPLI) was introduced in Stam and van Straaten (2012), and is defined as:

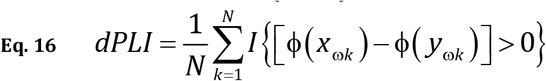

where “*I*” is the indicator function (equivalent to the Heaviside function), returning the value one if the argument is true, zero otherwise.

Stam and van Straaten (2012) assert that:

1. “x leads y” if dPLI is greater than 0.5.
2. “y leads x” if dPLI is smaller than 0.5.

An equivalent definition for the dPLI is:

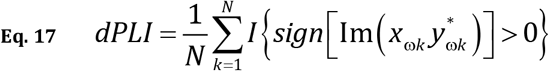

Note that:

1. The dPLI is invariant to mixing, as seen by plugging Eq. 7 into Eq. 17, under the condition that the mixing matrix be a positive definite matrix with positive determinant.
2. If one time series, say “y”, is multiplied by −1, then the dPLI will reverse and “go against the flow”. This property invalidates the dPLI as an estimator of direction of flow, since it violates the basic definition of Granger causality. This fact holds even for the ideal case of unmixed signals.

For convenience, the statistical significance tests will be based on the centered dPLI, denoted as CdPLI. It is defined as:

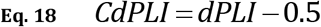

with the Stam and van Straaten (2012) assertions now written as:

1. “x leads y” if CdPLI is positive.
2. “y leads x” if CdPLI is negative.

### 4.5. The lagged coherence (LagCoh)

The lagged coherence was introduced in (Eq. 22 in Pascual-Marqui 2007a; Eqs. 29 and 31 in Pascual-Marqui 2007b; Eqs. 3.13, 3.14, and 3.17 in Pascual-Marqui et al 2011b). Its derivation is based on Geweke’s (1982) equations that express the total connectivity in *F*-transformed space as the sum of instantaneous and causal components. For time series filtered to a single discrete frequency, the total and instantaneous connectivities have clear, non-ambiguous definitions. This was used to derive the lagged connectivity in *F*-transformed space, which is trivially transformed into coherence-type measures. All the technical details can be found in the original lagged coherence papers quoted above.

Lagged coherence is defined as:

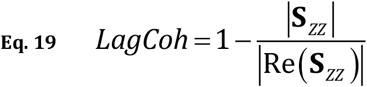

where Re(•) returns the real part of the complex valued argument, and 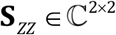is the Hermitian variance-covariance matrix for the observed time series. Plugging Eq. 5 into Eq. 19 gives:

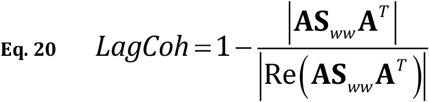

where *A* is the positive definite mixing matrix, and 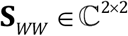is the Hermitian variance-covariance
matrix for the true, non-mixed, unobserved time series. This measure is obviously mixing-invariant because:

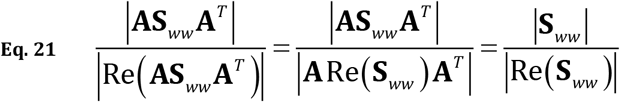

Thus, the LagCoh can be equivalently written as:

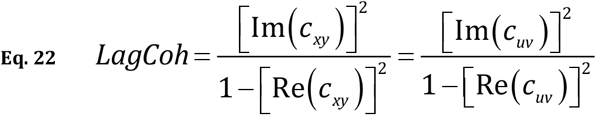

where*c_uv_* and *c_xy_* denote the complex valued coherence in the unmixed, and mixed cases, respectively.

### 4.6. The standardized imaginary covariance (sImCov)

The standardized imaginary covariance (sImCov) is defined as the t-statistic for the test with null hypothesis:

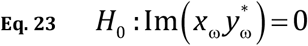

where “x” and “y” correspond to the observed, mixed signals.

The formal definition is:

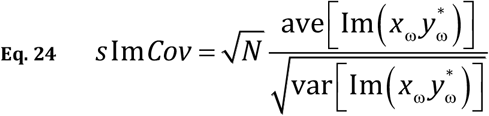

with:

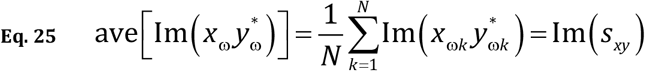

and:

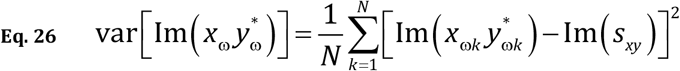

Plugging Eq. 7 into Eq. 24, Eq. 25, and Eq. 26 shows that the sImCov measure is mixing-invariant, i.e.:

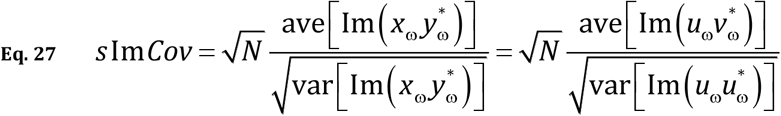

This new measure has the appeal of being a simple t-statistic, that has a corresponding p-value given by the t-distribution, that can be for direct statistical inference of the significance of the electrophysiological connection.

### 5. Simulations for a simple unidirectional causal model and its mixture

As before, let *u_tk_* and *v_tk_* denote the true, unobserved, unmixed signals, for *k* = 1…*N* trials, for *t* = 1…*N*_T_ time samples in each trial. And let *u*_ω*k*_ and *v*_ω*k*_ denote the complex valued discrete Fourier transform coefficients at discrete frequency ω.

Consider the model:

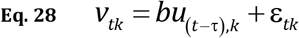

In this model, *u* and *ε* are independent, *ε* is the innovations time series, and “*u* is causing *v*” in the sense of Granger if τ>0, regardless of the sign of the regression coefficient *b*.

The discrete frequency domain model is:

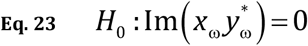

A very simple mixture is used in these simulations, namely:

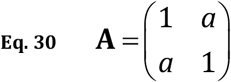

which is positive definite if −1 < *a* <+1. This mixture matrix produces the mixed variables:

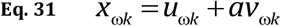

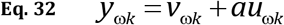

Simulated data is generated as follows:

1a. Given values for *a, b, N_τ_*, ω, *τ_0_* and *τ_J_*.
2a. For *k* = 1…*N* trials, generate the data in steps 3a to 5a:
3a. Generate random *u*_ω*k*_ and *ε*_ω*k*_. as independent complex normal variables (see e.g. Brillinger 2001), with zero mean and unit variance.
4a. Generate random time delay jitter as τ = τ_0_ + *U* (τ*_J_*), where *U* (τ*_J_*) returns a random uniformly distributed integer value in the range (-τ*_J_*)…(+τ*_J_*).
5a. Compute *v*_ω*k*_,*x*_ω*k*_, and *y*_ω*k*_.
6a. The data generated above consists of a sample of size *N*, for *k* =1…*N*, for the true non-mixed signals (*u*_ω*k*_,*v*_ω*k*_), and for the corresponding mixed signals (*x*_ω*k*_,*y*_ω*k*_). All connectivity measures (ImCoh, LagCoh, PLI, wPLI, CdPLI, sImCov) are then computed, using the respective equations described in earlier sections of this paper. Additionally, two tailed significance tests for zero connectivity are performed via non-parametric randomization for regression (based on 1000 randomizations each), as described in Manly (2006), at levels p<0.05, 0.1, and 0.2. Furthermore, the two tailed t-distribution probability for the sImCov as a t-statistic is also calculated.

The above procedure, steps 1a to 6a, are repeated 1000 times. For each connectivity measure, out of 1000 tests, the number of true positive detections for significant connectivity at levels p<0.05,0.1, and 0.2 gives the rate of true positive detection of the measure.

## 6. Results and discussion

The results, corresponding to true positive detection rates in percent (0…100), are shown in Table 1 (a,b,c) for the mixed signals, and in Table 2 (a,b,c) for the original unmixed signals. The parameters are *a, b, N*, ω, τ_0_ and τ*_J_* are indicated for each simulation. In all cases, *N_T_* = 128 was used.

**Table 1a:** True positive detection rates for all connectivity measures at ***p<0.05*** (first seven columns) for ***mixture*** of time series. First six columns are based on randomization tests for 1000 realizations, and seventh column is based on the t-distribution. Highlighted in light yellow is the best connectivity measure. Columns 8–13 define the parameters used in the simulation.

| ImCoh | LagCoh | PLI | wPLI | CdPLI | slmCov | slmCov(t) | #Trials | Freq | Tau | Tau(jitter) | a | RegCoeff |
| --- | --- | --- | --- | --- | --- | --- | --- | --- | --- | --- | --- | --- |
| 0 | 10.7 | 5.2 | 8 | 5.2 | 8.9 | 8 | 20 | 2 | 1 | 0.00E+00 | -8.00E-01 | -1.00E+00 |
| 0 | 48 | 19 | 38.8 | 19 | 36.6 | 35 | 20 | 6 | 1 | 0.00E+00 | -8.00E-01 | -1.00E+00 |
| 0 | 89.5 | 51.1 | 84.2 | 51.1 | 80.5 | 79.7 | 20 | 11 | 1 | 0.00E+00 | -8.00E-01 | -1.00E+00 |
| 0 | 99.2 | 79.8 | 98.2 | 79.8 | 97.9 | 97.8 | 20 | 18 | 1 | 0.00E+00 | -8.00E-01 | -1.00E+00 |
| 0 | 100 | 90.7 | 100 | 90.7 | 99.6 | 99.7 | 20 | 25 | 1 | 0.00E+00 | -8.00E-01 | -1.00E+00 |
| 0 | 7.8 | 4.8 | 6.2 | 4.8 | 6.5 | 6.4 | 20 | 6 | 1 | 0.00E+00 | 8.00E-01 | 1.00E-01 |
| 0 | 9.4 | 5.6 | 7.9 | 5.6 | 7.6 | 6.6 | 20 | 11 | 1 | 0.00E+00 | 8.00E-01 | 1.00E-01 |
| 0 | 10.8 | 6.7 | 10.2 | 6.7 | 9.5 | 8.8 | 20 | 18 | 1 | 0.00E+00 | 8.00E-01 | 1.00E-01 |
| 0 | 12.4 | 6.8 | 11 | 6.8 | 10.8 | 9.8 | 20 | 25 | 1 | 0.00E+00 | 8.00E-01 | 1.00E-01 |
| 0 | 7.9 | 5.2 | 5.8 | 5.2 | 5.9 | 5.7 | 20 | 2 | 2 | 3.00E+00 | 8.00E-01 | 1.00E-01 |
| 0 | 9.1 | 5.7 | 7.5 | 5.7 | 7.1 | 6.8 | 20 | 6 | 2 | 3.00E+00 | 8.00E-01 | 1.00E-01 |
| 0 | 9.6 | 5.4 | 8.2 | 5.4 | 7.7 | 7 | 20 | 11 | 2 | 3.00E+00 | 8.00E-01 | 1.00E-01 |
| 0 | 7.7 | 4.3 | 6.1 | 4.3 | 6 | 5.5 | 20 | 18 | 2 | 3.00E+00 | 8.00E-01 | 1.00E-01 |
| 0 | 6.7 | 4.6 | 5.5 | 4.6 | 5.5 | 4.8 | 20 | 25 | 2 | 3.00E+00 | 8.00E-01 | 1.00E-01 |
| 0 | 7.4 | 5.6 | 7.7 | 5.6 | 7.6 | 7.1 | 200 | 2 | 2 | 3.00E+00 | 8.00E-01 | 1.00E-01 |
| 0 | 16.8 | 9 | 15.9 | 9 | 16.4 | 16.2 | 200 | 6 | 2 | 3.00E+00 | 8.00E-01 | 1.00E-01 |
| 0 | 19.8 | 10.3 | 19.3 | 10.3 | 19.8 | 18.8 | 200 | 11 | 2 | 3.00E+00 | 8.00E-01 | 1.00E-01 |
| 0 | 6.9 | 5.8 | 7 | 5.8 | 7.3 | 6.8 | 200 | 18 | 2 | 3.00E+00 | 8.00E-01 | 1.00E-01 |
| 0 | 5.9 | 5.9 | 5.8 | 5.9 | 5.5 | 5.3 | 200 | 25 | 2 | 3.00E+00 | 8.00E-01 | 1.00E-01 |

**Table 1b:** True positive detection rates for all connectivity measures at ***p<0.1*** (first seven columns) for *mixture* of time series. First six columns are based on randomization tests for 1000 realizations, and seventh column is based on the t-distribution. Highlighted in dark yellow is the best connectivity measure. Columns 8–13 define the parameters used in the simulation.

| ImCoh | LagCoh | PLI | wPLI | CdPLI | slmCov | slmCov(t) | #Trials | Freq | Tau | Tau(jitter) | a | RegCoeff |
| --- | --- | --- | --- | --- | --- | --- | --- | --- | --- | --- | --- | --- |
| 0 | 19.1 | 8.9 | 15.3 | 8.9 | 15.7 | 15.4 | 20 | 2 | 1 | 0.00E+00 | -8.00E-01 | -1.00E+00 |
| 0 | 59.2 | 23.3 | 52.8 | 23.3 | 51.8 | 50.8 | 20 | 6 | 1 | 0.00E+00 | -8.00E-01 | -1.00E+00 |
| 0 | 93.9 | 56.3 | 90.8 | 56.3 | 89.6 | 90 | 20 | 11 | 1 | 0.00E+00 | -8.00E-01 | -1.00E+00 |
| 0 | 99.9 | 84.2 | 99.5 | 84.2 | 99.3 | 99.3 | 20 | 18 | 1 | 0.00E+00 | -8.00E-01 | -1.00E+00 |
| 0 | 100 | 92.8 | 100 | 92.8 | 100 | 100 | 20 | 25 | 1 | 0.00E+00 | -8.00E-01 | -1.00E+00 |
| 0 | 12.9 | 8.1 | 12.2 | 8.1 | 12.1 | 11.6 | 20 | 6 | 1 | 0.00E+00 | 8.00E-01 | 1.00E-01 |
| 0 | 15 | 8.4 | 13.1 | 8.4 | 13.7 | 12.9 | 20 | 11 | 1 | 0.00E+00 | 8.00E-01 | 1.00E-01 |
| 0 | 17.4 | 9.5 | 15.3 | 9.5 | 14.7 | 15 | 20 | 18 | 1 | 0.00E+00 | 8.00E-01 | 1.00E-01 |
| 0 | 19.4 | 9.4 | 17.4 | 9.4 | 16.9 | 16.7 | 20 | 25 | 1 | 0.00E+00 | 8.00E-01 | 1.00E-01 |
| 0 | 12.7 | 8.6 | 11.5 | 8.6 | 11.9 | 11.5 | 20 | 2 | 2 | 3.00E+00 | 8.00E-01 | 1.00E-01 |
| 0 | 13.9 | 8.3 | 12.7 | 8.3 | 13 | 12.6 | 20 | 6 | 2 | 3.00E+00 | 8.00E-01 | 1.00E-01 |
| 0 | 14.5 | 8.5 | 13.4 | 8.5 | 13.4 | 12.9 | 20 | 11 | 2 | 3.00E+00 | 8.00E-01 | 1.00E-01 |
| 0 | 12.4 | 7.1 | 11.9 | 7.1 | 12.1 | 11 | 20 | 18 | 2 | 3.00E+00 | 8.00E-01 | 1.00E-01 |
| 0 | 12.9 | 6.7 | 11.8 | 6.7 | 11.6 | 11.1 | 20 | 25 | 2 | 3.00E+00 | 8.00E-01 | 1.00E-01 |
| 0 | 13.2 | 11.2 | 13.4 | 11.2 | 13 | 13.1 | 200 | 2 | 2 | 3.00E+00 | 8.00E-01 | 1.00E-01 |
| 0 | 25.9 | 14.4 | 25 | 14.4 | 24.8 | 25.2 | 200 | 6 | 2 | 3.00E+00 | 8.00E-01 | 1.00E-01 |
| 0 | 29.4 | 16.1 | 28.6 | 16.1 | 28.4 | 28.8 | 200 | 11 | 2 | 3.00E+00 | 8.00E-01 | 1.00E-01 |
| 0 | 12.9 | 10.5 | 12.5 | 10.5 | 12.5 | 12.8 | 200 | 18 | 2 | 3.00E+00 | 8.00E-01 | 1.00E-01 |
| 0 | 11.4 | 10.6 | 11.5 | 10.6 | 11.4 | 11.9 | 200 | 25 | 2 | 3.00E+00 | 8.00E-01 | 1.00E-01 |

**Table 1c:** True positive detection rates for all connectivity measures at ***p<0.2*** (first seven columns) for ***mixture*** of time series. First six columns are based on randomization tests for 1000 realizations, and seventh column is based on the t-distribution. Highlighted in green is the best connectivity measure. Columns 8–13 define the parameters used in the simulation.

| ImCoh | LagCoh | PLI | wPLI | CdPLI | slmCov | slmCov(t) | #Trials | Freq | Tau | Tau(jitter) | a | RegCoeff |
| --- | --- | --- | --- | --- | --- | --- | --- | --- | --- | --- | --- | --- |
| 0 | 31.8 | 16 | 28.5 | 22.4 | 27.1 | 27.8 | 20 | 2 | 1 | 0.00E+00 | -8.00E-01 | -1.00E+00 |
| 0 | 71.7 | 37.2 | 68.1 | 49 | 67.1 | 67.5 | 20 | 6 | 1 | 0.00E+00 | -8.00E-01 | -1.00E+00 |
| 0 | 96.6 | 69.4 | 95.8 | 79.5 | 95.8 | 96 | 20 | 11 | 1 | 0.00E+00 | -8.00E-01 | -1.00E+00 |
| 0 | 100 | 92.1 | 100 | 95.5 | 99.9 | 99.9 | 20 | 18 | 1 | 0.00E+00 | -8.00E-01 | -1.00E+00 |
| 0.3 | 100 | 98 | 100 | 99.1 | 100 | 100 | 20 | 25 | 1 | 0.00E+00 | -8.00E-01 | -1.00E+00 |
| 0 | 23.4 | 13.5 | 21.7 | 17.3 | 21.3 | 21.1 | 20 | 6 | 1 | 0.00E+00 | 8.00E-01 | 1.00E-01 |
| 0 | 24.9 | 13.7 | 23.6 | 16.7 | 23.2 | 23.1 | 20 | 11 | 1 | 0.00E+00 | 8.00E-01 | 1.00E-01 |
| 0 | 27.8 | 15.4 | 26 | 18.1 | 25.7 | 25.7 | 20 | 18 | 1 | 0.00E+00 | 8.00E-01 | 1.00E-01 |
| 0 | 30.2 | 16.2 | 27.8 | 18.8 | 27.8 | 28.2 | 20 | 25 | 1 | 0.00E+00 | 8.00E-01 | 1.00E-01 |
| 0 | 23.3 | 13 | 20.9 | 17.2 | 20.7 | 21.3 | 20 | 2 | 2 | 3.00E+00 | 8.00E-01 | 1.00E-01 |
| 0 | 24.4 | 13.8 | 22.3 | 17.2 | 22.6 | 22.3 | 20 | 6 | 2 | 3.00E+00 | 8.00E-01 | 1.00E-01 |
| 0 | 25.2 | 14.3 | 23.2 | 17.7 | 23.3 | 23 | 20 | 11 | 2 | 3.00E+00 | 8.00E-01 | 1.00E-01 |
| 0 | 23.4 | 12.1 | 21.2 | 17.3 | 20.8 | 21 | 20 | 18 | 2 | 3.00E+00 | 8.00E-01 | 1.00E-01 |
| 0 | 23 | 12.4 | 20.5 | 18.3 | 21.1 | 21.3 | 20 | 25 | 2 | 3.00E+00 | 8.00E-01 | 1.00E-01 |
| 0 | 24.4 | 19.1 | 24 | 21 | 23.9 | 23.8 | 200 | 2 | 2 | 3.00E+00 | 8.00E-01 | 1.00E-01 |
| 0 | 38.6 | 26.1 | 38.7 | 27.8 | 38.7 | 38.6 | 200 | 6 | 2 | 3.00E+00 | 8.00E-01 | 1.00E-01 |
| 0 | 42.7 | 29.2 | 42.2 | 30.4 | 42.2 | 42.7 | 200 | 11 | 2 | 3.00E+00 | 8.00E-01 | 1.00E-01 |
| 0 | 23.5 | 19.5 | 22.9 | 20.4 | 22.8 | 23.2 | 200 | 18 | 2 | 3.00E+00 | 8.00E-01 | 1.00E-01 |
| 0 | 22.8 | 19.3 | 22.5 | 19.8 | 22.6 | 21.6 | 200 | 25 | 2 | 3.00E+00 | 8.00E-01 | 1.00E-01 |

**Table 2a:** True positive detection rates for all connectivity measures at ***p<0.05*** (first seven columns) for the original ***unmixed*** signals. First six columns are based on randomization tests for 1000 realizations, and seventh column is based on the t-distribution. Highlighted in light yellow is the best connectivity measure. Columns 8–13 define the parameters used in the simulation.

| ImCoh | LagCoh | PLI | wPLI | CdPLI | slmCov | slmCov(t) | #Trials | Freq | Tau | Tau(jitter) | a | RegCoeff |
| --- | --- | --- | --- | --- | --- | --- | --- | --- | --- | --- | --- | --- |
| 1.3 | 8.2 | 5.3 | 7.2 | 5.3 | 8.4 | 8 | 20 | 2 | 1 | 0.00E+00 | 0 | -1.00E+00 |
| 17.4 | 44.7 | 18 | 38.7 | 18 | 35.6 | 35 | 20 | 6 | 1 | 0.00E+00 | 0 | -1.00E+00 |
| 70.3 | 87.1 | 49.3 | 83 | 49.3 | 77.9 | 79.7 | 20 | 11 | 1 | 0.00E+00 | 0 | -1.00E+00 |
| 98.2 | 98.9 | 78.9 | 98.1 | 78.9 | 97.6 | 97.8 | 20 | 18 | 1 | 0.00E+00 | 0 | -1.00E+00 |
| 100 | 100 | 90.3 | 99.9 | 90.3 | 99.4 | 99.7 | 20 | 25 | 1 | 0.00E+00 | 0 | -1.00E+00 |
| 6.2 | 6.2 | 4.9 | 6.7 | 4.9 | 6.3 | 6.4 | 20 | 6 | 1 | 0.00E+00 | 0 | 1.00E-01 |
| 7.4 | 8 | 5.4 | 8.1 | 5.4 | 6.8 | 6.6 | 20 | 11 | 1 | 0.00E+00 | 0 | 1.00E-01 |
| 9 | 9.1 | 6.6 | 8.8 | 6.6 | 8.3 | 8.8 | 20 | 18 | 1 | 0.00E+00 | 0 | 1.00E-01 |
| 10.1 | 10.3 | 6.6 | 10.2 | 6.6 | 9.5 | 9.8 | 20 | 25 | 1 | 0.00E+00 | 0 | 1.00E-01 |
| 5.3 | 5.8 | 5.2 | 6.5 | 5.2 | 5.9 | 5.7 | 20 | 2 | 2 | 3.00E+00 | 0 | 1.00E-01 |
| 7.5 | 7.8 | 6 | 7.5 | 6 | 6.3 | 6.8 | 20 | 6 | 2 | 3.00E+00 | 0 | 1.00E-01 |
| 8.1 | 7.9 | 5.6 | 7.5 | 5.6 | 6.9 | 7 | 20 | 11 | 2 | 3.00E+00 | 0 | 1.00E-01 |
| 5.9 | 5.6 | 4.4 | 5.6 | 4.4 | 5.5 | 5.5 | 20 | 18 | 2 | 3.00E+00 | 0 | 1.00E-01 |
| 4.9 | 4.7 | 4.5 | 4.7 | 4.5 | 4.9 | 4.8 | 20 | 25 | 2 | 3.00E+00 | 0 | 1.00E-01 |
| 6.9 | 7 | 5.5 | 7.2 | 5.5 | 7.5 | 7.1 | 200 | 2 | 2 | 3.00E+00 | 0 | 1.00E-01 |
| 16.4 | 16.4 | 8.9 | 16.5 | 8.9 | 16.5 | 16.2 | 200 | 6 | 2 | 3.00E+00 | 0 | 1.00E-01 |
| 19.5 | 19.5 | 10 | 19.6 | 10 | 19.2 | 18.8 | 200 | 11 | 2 | 3.00E+00 | 0 | 1.00E-01 |
| 6.6 | 6.6 | 5.5 | 6.8 | 5.5 | 6.8 | 6.8 | 200 | 18 | 2 | 3.00E+00 | 0 | 1.00E-01 |
| 5.8 | 5.8 | 6 | 5.5 | 6 | 5.5 | 5.3 | 200 | 25 | 2 | 3.00E+00 | 0 | 1.00E-01 |

**Table 2b:** True positive detection rates for all connectivity measures at ***p<0.1***(first seven columns) for the original ***unmixed*** signals. First six columns are based on randomization tests for 1000 realizations, and seventh column is based on the t-distribution. Highlighted in dark yellow is the best connectivity measure. Columns 8–13 define the parameters used _ in the simulation.

| ImCoh | LagCoh | PLI | wPLI | CdPLI | slmCov | slmCov(t) | #Trials | Freq | Tau | Tau(jitter) | a | RegCoeff |
| --- | --- | --- | --- | --- | --- | --- | --- | --- | --- | --- | --- | --- |
| 3.7 | 17.1 | 7.3 | 15.4 | 7.3 | 14.6 | 15.4 | 20 | 2 | 1 | 0.00E+00 | 0 | -1.00E+00 |
| 31.7 | 56.9 | 22.1 | 52.6 | 22.1 | 50.3 | 50.8 | 20 | 6 | 1 | 0.00E+00 | 0 | -1.00E+00 |
| 83.4 | 92.8 | 55.5 | 90.3 | 55.5 | 89.1 | 90 | 20 | 11 | 1 | 0.00E+00 | 0 | -1.00E+00 |
| 99.5 | 99.7 | 83.4 | 99.5 | 83.4 | 99.1 | 99.3 | 20 | 18 | 1 | 0.00E+00 | 0 | -1.00E+00 |
| 100 | 100 | 92.4 | 100 | 92.4 | 100 | 100 | 20 | 25 | 1 | 0.00E+00 | 0 | -1.00E+00 |
| 11.4 | 11.2 | 6.3 | 11.4 | 6.3 | 11.2 | 11.6 | 20 | 6 | 1 | 0.00E+00 | 0 | 1.00E-01 |
| 12.5 | 12.2 | 6.8 | 12.9 | 6.8 | 12.3 | 12.9 | 20 | 11 | 1 | 0.00E+00 | 0 | 1.00E-01 |
| 13.8 | 14.1 | 8.4 | 15.1 | 8.4 | 13.9 | 15 | 20 | 18 | 1 | 0.00E+00 | 0 | 1.00E-01 |
| 15.6 | 15.5 | 8.5 | 16 | 8.5 | 16.5 | 16.7 | 20 | 25 | 1 | 0.00E+00 | 0 | 1.00E-01 |
| 11 | 11.1 | 6.6 | 10.7 | 6.6 | 10.8 | 11.5 | 20 | 2 | 2 | 3.00E+00 | 0 | 1.00E-01 |
| 12.1 | 11.7 | 7.2 | 12.4 | 7.2 | 12 | 12.6 | 20 | 6 | 2 | 3.00E+00 | 0 | 1.00E-01 |
| 12 | 12.2 | 7 | 12.7 | 7 | 12.7 | 12.9 | 20 | 11 | 2 | 3.00E+00 | 0 | 1.00E-01 |
| 10.4 | 10.4 | 5.9 | 10.6 | 5.9 | 10.6 | 11 | 20 | 18 | 2 | 3.00E+00 | 0 | 1.00E-01 |
| 9.7 | 10.2 | 5.5 | 10.6 | 5.5 | 10.6 | 11.1 | 20 | 25 | 2 | 3.00E+00 | 0 | 1.00E-01 |
| 12.6 | 12.7 | 10.5 | 12.9 | 10.5 | 13 | 13.1 | 200 | 2 | 2 | 3.00E+00 | 0 | 1.00E-01 |
| 25.3 | 25.4 | 14.5 | 25.5 | 14.5 | 25.3 | 25.2 | 200 | 6 | 2 | 3.00E+00 | 0 | 1.00E-01 |
| 29.5 | 29.5 | 15.4 | 28.4 | 15.4 | 28.5 | 28.8 | 200 | 11 | 2 | 3.00E+00 | 0 | 1.00E-01 |
| 12 | 12.1 | 10.6 | 12.2 | 10.6 | 12.5 | 12.8 | 200 | 18 | 2 | 3.00E+00 | 0 | 1.00E-01 |
| 11.4 | 11.3 | 10.7 | 11.2 | 10.7 | 11.7 | 11.9 | 200 | 25 | 2 | 3.00E+00 | 0 | 1.00E-01 |

**Table 2c:**
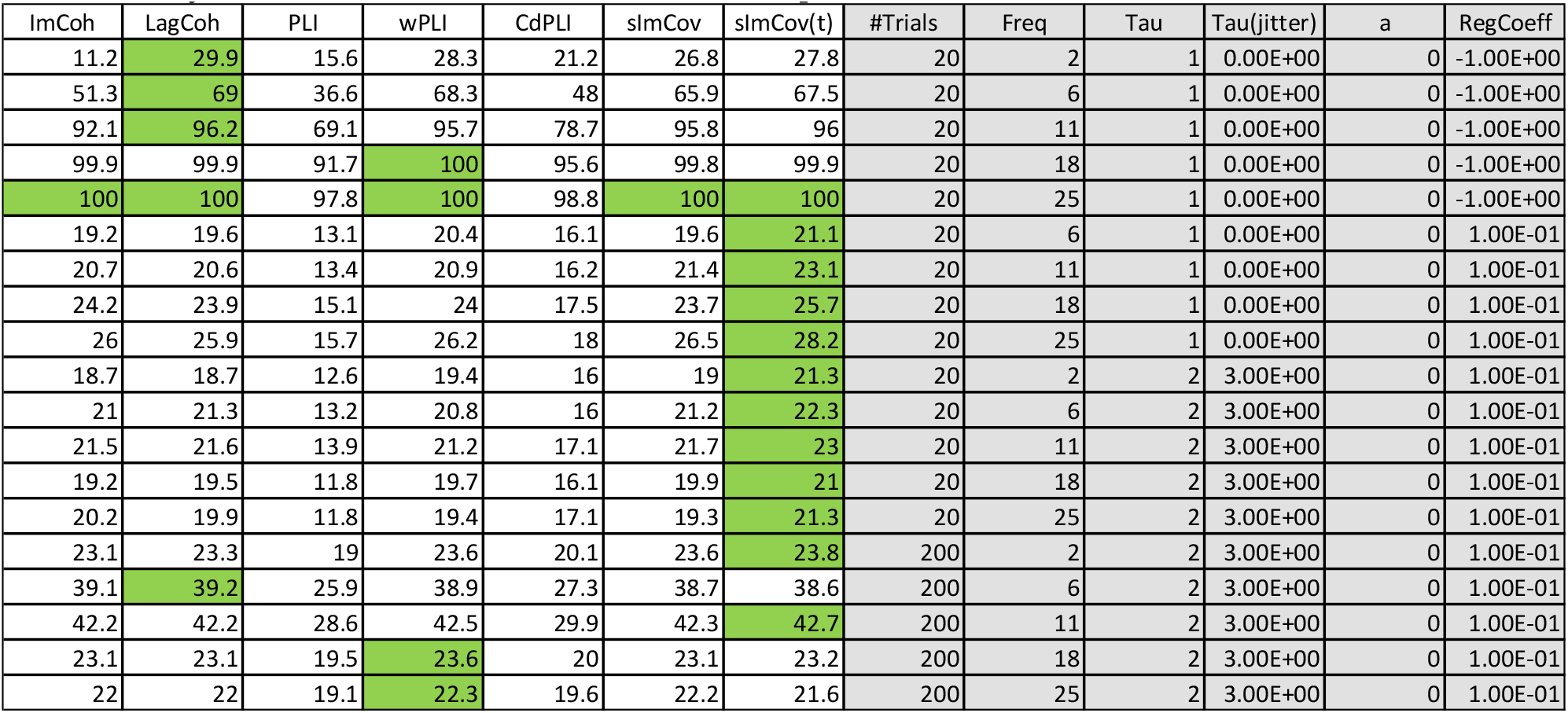
True positive detection rates for all connectivity measures at ***p<0.2*** (first seven columns) for the original ***unmixed*** signals. First six columns are based on randomization tests for 1000 realizations, and seventh column is based on the t-distribution. Highlighted in green is the best connectivity measure. Columns 8–13 define the parameters used in the simulation.

The results show that in the majority of cases, for the mixed signals, the LagCoh is the clear best electrophysiological connectivity measure. As for the ideal case of the unmixed signals, the LagCoh and the sImCov in its theoretical form (as a t-statistic, with probability calculated from the t-distribution) share the position of best electrophysiological connectivity measure.

These results are very limited in scope:

1. The model is very simple. Other models must be studied.
2. The actual number of simulations are is small. More exhaustive exploration of all parameters is needed.

Note that all statistical tests presented here correspond to what is termed “subject level statistics”. Since LagCoh has the best performance under the limited conditions tested here, it can be safely assumed that it will very likely have the best performance in group level statistics, where LagCoh is calculated for groups of subjects, under different conditions. See e.g. (Vanneste et al (2011), which shows this type of analysis.

Finally, the following facts deserve special emphasis:

1. PLI and dPLI have very similar performance.
2. Under the very limited conditions tested here, the dPLI has a rather poor performance, as compared to LagCoh, sImCov, and wPLI.
3. The dPLI is an invalid estimator of flow direction, i.e. it reverses and “goes against the flow” by merely changing the sign of one of the time series, a fact that violates the basic definition of Granger causality. This occurs in the dPLI estimator for the model in Eq. 29 by changing the regression coefficient from “+*b*” to “-*b*”, for any non-zero “*b*” value. Whereas the concept of Granger causality asserts that the direction of causality in the model considered here is “x causes y for τ>0”, dPLI reverses the direction if “b<0”. Note that the sign of a time series can be completely arbitrary. For instance, consider ECoG recordings at two well separated cortical sites, and where at each site, a local bipolar recording is available which guarantees a very local electric potential gradient measurement at each site. But the signs of the bipolar recordings are arbitrary and immaterial for the purpose of estimating the direction of flow of information in the sense of Granger causality. In this practical example, dPLI will give an arbitrary direction, depending on the signs of the signals.

